# Mosaic chromosomal aneuploidy detection by sequencing (MAD-seq)

**DOI:** 10.1101/142299

**Authors:** Yu Kong, Esther R. Berko, Anthony Marcketta, Shahina B. Maqbool, Claudia A. Simões-Pires, David F. Kronn, Kenny Q. Ye, Masako Suzuki, Adam Auton, John M. Greally

## Abstract

Current approaches to detect and characterize mosaic chromosomal aneuploidy are limited by sensitivity, efficiency, cost or the need to culture cells. We describe a combination of a new sequencing-based assay and a novel analytical approach that allows low levels of mosaicism for chromosomal aneuploidy to be detected, assigned to a meiotic or mitotic origin, and quantified as a proportion of the cells in the sample. We show results from a multi-ethnic assay design that is suitable for populations of diverse racial and ethnic origins, and how the *MADSEQ* analytical approach applied to exome sequencing data reveals unrecognized aneuploidy in 1000 Genomes samples and cell lines from public repositories. We have made the assay design and analytical software open for unrestricted use, with the goal that it can be applied in clinical samples to allow new insights into the unrecognized prevalence of mosaic chromosomal aneuploidy and its phenotypic associations.

## INTRODUCTION

Somatic mosaicism occurs when a single population of cells acquires a subpopulation with a different genotype. While mosaicism is an unavoidable consequence of the low level of background mutation, making every multicellular organism a mosaic to some degree, the pathogenic consequences of mosaicism are most apparent when (a) it occurs early in development, and therefore affects a substantial proportion of cells forming one or more organs, and (b) the genotypic alteration is a detrimental mutation.

Chromosomal abnormalities are found at surprisingly high rates in human zygotes, with estimates that as many as three-quarters of these early embryos contain aneuploid cells.^1,2^ It has been assumed that there is a selective growth and survival advantage for any subset of normal diploid cells contained within these embryos,^3^ accounting for the generally phenotypically and chromosomally normal outcomes observed. It remains possible, however, that some of the aneuploid cells present in the zygote persist through development. This is recognized more frequently in placental than in embryonic tissues. When defined by the presence of aneuploidy in chorionic villus sampling (CVS) samples from the placenta, and the failure to detect such aneuploid cells in fetal amniocytes or cells from the newborn, it is referred to as confined placental mosaicism (CPM).^4^ With current technologies, approximately 0.8–2.0% of all CVS speciments are found to have mosaic aneuploidy, and a subset of 10-20% of “confined” placental mosaicism is now recognized to have the same mosaic aneuploidy in the fetus.^5–10^ CPM has been found at higher rates (up to ~15%^11^) in cases of intra-uterine growth restriction (IUGR).

The prevalence of mosaic chromosomal aneuploidy is not known to be substantial in humans, but it is almost certainly under-recognized. The nature of the developmental event is such that it may only affect an anatomically-restricted group of cells in the body, whereas routine genetic testing is almost always performed upon DNA from peripheral blood leukocytes, looking for constitutive, germ line mutations. Blood, in particular when cultured, is especially poor as a choice of cells for detection of mosaic aneuploidy (exemplified by the failure to detect tetrasomy 12p in Pallister-Killian syndrome^12–14^), with blood-derived lymphoblastoid cell lines (LCLs) likely to be even worse, given their striking oligoclonality.^15^ Current large-scale studies of human phenotypes, mostly based on blood or LCL DNA, are therefore unlikely to be optimally sensitive for detection of the presence of mosaic chromosomal aneuploidy. It follows that even the studies to date involving thorough analyses of molecular genomic data are likely to have systematically missed evidence for mosaic aneuploidy occurring in human subjects. In those studies in which mosaic chromosomal aneuploidy was specifically sought, it was found to be associated with certain phenotypes, including several reports of mosaicism for chromosomal aneuploidy in peripheral blood in children with autism spectrum disorder (ASD).^16–18^ Given the limitations of using blood to detect aneuploid cells, it is likely that mosaic chromosomal aneuploidy in ASD is not limited to these specific reported individuals but is more prevalent.

Current technologies to detect and quantify mosaic chromosomal aneuploidy include karyotyping of chromosomes using large numbers of metaphase cells generated using cell culture,^19^ fluorescence *in situ* hybridization (FISH) using probes detecting the aneuploid chromosome,^20^ microarrays to genotype the sample and measure the relative fluorescence of minor alleles,^21^ single cell sequencing,^22^ and whole genome sequencing.^23^ These approaches all have their relative strengths, but we currently lack an assay that combines relative ease and cost-effectiveness, sensitive detection of low proportions of aneuploid cells, no requirement for cell culture, characterization of the original mitotic or meiotic origin of the mutation, suitability for multiple racial and ethnic populations, and a supporting analytical software resource. To address this need, we developed the MAD-seq assay and its supporting, open source *MADSEQ* analytical software package.

## MATERIALS AND METHODS

### Molecular Assays

#### Cell line mixing experiments

DNA extracted from two individuals’ LCLs (GM06990, CEU, female and GM19239, YRI, male) was mixed at different proportions, choosing 100%:0%, 99.5%:0.5%, 95%:5%, 90%:10%, 75%:25%, 50%:50% to mimic different levels of mosaicism. The DNA was sequenced following a capture protocol using the SeqCap EZ Choice system from Roche-NimbleGen. The list of targeted regions and probes used for the v1MAD-seq design can be found in our dbGaP submission. Knowing the genotype of both samples, we were able to extract sites that mimic the different types of mosaic aneuploidy. Specifically, to simulate mitotic aneuploidy, we first extracted loci with different genotypes in the two cell lines, 0/1 in CEU and 1/1 in YRI to mimic over-representation of the alternative allele, and 0/1 in CEU and 0/0 in YRI to mimic under-representation of the alternative allele. The two mixtures separated further when a higher proportion of YRI DNA was mixed with CEU DNA (**Table S1**). Because these mixtures of DNA alter the distribution of alternative allele frequency without changing the actual copy number of chromosomes, we applied the model without the coverage module.

#### Exome sequencing of a patient with hemihyperplasia

DNA was extracted from fibroblasts cultured from skin biopsies from the affected and unaffected sides of the body of a patient with hemihyperplasia (OMIM 235000). We performed exome sequencing of the sample from the affected side of this patient using the SeqCap EZ Exome Enrichment Kit v3.0 (Roche-NimbleGen) and 100 base paired-end sequencing on the Illumina HiSeq 2500 system. The average coverage was 142.1 X.

#### Targeted sequencing

DNA was purchased from Coriell for the samples HG01939, NA00682 and NA01454, and DNA was extracted from fibroblasts (AG13074, GM00496 and GM00503) and LCLs (GM06990, GM19239), and buccal epithelial cells of one patient (F44P110), and a human embryonic cell line (H1). DNA extracted from the normal and abnormal cultured fibroblasts of the hemihypertrophy patient was also included. We used the Roche NimbleGen SeqCap EZ Choice system to capture the multi-ethnic design of the 105,703 common SNPs described below. All of the samples were sequenced with 100 bp paired-end sequencing using the Illumina HiSeq 2500 (Illumina, San Diego, CA), generating an average coverage of 134.6X.

#### Software availability

The *MADSEQ* model was implemented and released as a Bioconductor R package. The source code and instructions to use this package are available at http://bioconductor.org/packages/MADSEQ/

#### Sequencing depth correction

G+C content can vary in the genome and influence the number of reads generated at each captured region, potentially introducing bias into aneuploidy detection. We therefore used LOESS correction to correct in our package for such bias. Given the targeted region and bam file, the average coverage for each targeted region (*raw_cov_i_*) is calculated by a coverage function from the *MADSEQ* R package called *GenomicAlignments*. If more than one sample is sequenced during the same capture protocol, quantile normalization is first applied to the coverage across all samples. The G+C content (*gc_i_*) for each targeted region is then calculated as the G+C percentage of the reference genome (excluding Ns). Coverage for each targeted region was grouped by 0.1% increments of G+C content, and the average coverage for each level of G+C content was calculated. The scatterplot representing the G+C content plotted against the average coverage for each G+C level can be produced as part of the *MADSEQ* pipeline (**Figure S10**). The regression curve between coverage and G+C content was fitted by LOESS. The GC content for the *i^th^* region is *gc_i_*, the fitted coverage for this region denoted as *cov_gc_i__*. The expected coverage (*cov_exp_*) is set to the median of read depth across all regions. The corrected coverage for the *i^th^* region (*corrected_cov_i_*) is then calculated as:

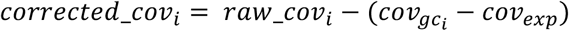

#### Bayesian models

Our statistical model for detecting mosaic aneuploidy consists of two parts. For each chromosome, we first consider the distribution of the alternative allele frequencies at the heterozygous sites; secondly, we consider the distribution of sequencing depth at these loci. If mosaic aneuploidy is present in the sample, we expect the distribution of both the alternative allele frequency and sequencing depth to deviate from that expected in a simple diploid sample. Here we describe each part of the model separately:

##### 1. Detection of aneuploidy from alternative allelic fractions (AAF)

The alternative allele fraction is the proportion of reads carrying the alternative allele at a given heterozygous site, calculated as the alternative allelic depth divided by the total read depth. If there is no aneuploidy in any of the sampled cells, then the AAFs at heterozygous sites are expected to be centered around 0.5 (ignoring confounding biases such as reference bias^36^). However, if a fraction of cells within the sample are aneuploid, then the distribution of AAFs will deviate from the expected midpoint, and instead be better described by a mixture of distributions, where the number and parameters of the mixture components depending on the origin of the aneuploidy and the degree of mosaic aneuploidy.

1. **Model_0_: diploid chromosome**. For a normal, diploid chromosome state, the AAF at heterozygous sites is expected to be a single distribution centered around the midpoint (average AAF across all heterozygous sites). In this situation, we model the alternative allelic depth (*AD_i_*) for biallelic heterozygous site *i* as a simple beta-binomial distribution, given the read depth for the *i^th^* site *N_i_*:

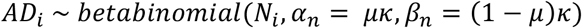 Here, *α* and *β* are determined by the prior *μ* and *κ. μ* denotes the midpoint, namely the average AAF across all the heterozygous sites. *κ* represents the variance of the AAF. We model *κ* as a gamma distribution:

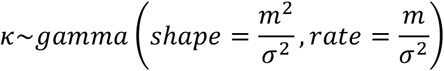 In our model, the larger the *κ* is, the smaller the variance for the beta distribution. For the purpose of Bayesian inference, we assigned the prior for this gamma distribution as *m* = 10, *σ* = 10 to represent a flat prior distribution. To account for noise normally present in high-throughput sequencing data, we added an additional outlier component weighted as 1% (*ω*_0_ = 0.01; *ω_n_* + *ω*_0_ = 1) of all heterozygous reads. The outlier component is modeled by a uniform beta-binomial distribution as

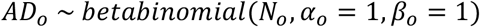 If there are K heterozygous sites from one chromosome, and mu and Kappa are constant for all n=1,…K, the likelihood of the data is then given by:

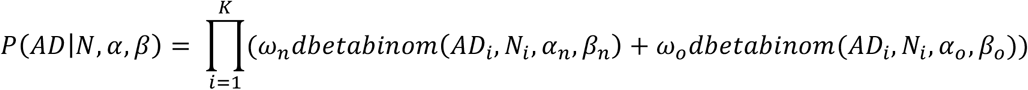

where *P*(*AD*|*N, α, β*) are vectors of the parameters.
2. **Model_1_: mosaic monosomy.** Mosaic monosomy is the consequence of the loss of one chromosome in a subset of cells. In mosaic monosomy, the AAF separates from the midpoint into 2 mixtures (Figure 2). One mixture is shifted toward lower values due to the over-representation of the reference allele, and the other shifted toward higher values due to the over-representation of the alternative allele. We assume that the two mixtures have equal weight (*ω*_1_ = *ω*_2_ = 0.495) and variance (*κ*), with the same outlier component (*ω*_0_ = 0.01) described in model_0_, the allelic depth of heterozygous sites can be modelled as:

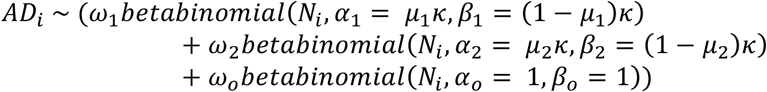 The more monosomic cells in the sample, the further these two mixtures will separate (**Figure S2**). The priors *μ*_1_ and *μ*_2_, which are the average AAF of the two separated mixtures, are determined by the fraction of aneuploidy cells, *f*. Given the expected midpoint (calculated as average AAF for all heterozygous sites from the whole genome) *m*, the expected mean AAF of the two mixtures (*μ*_1_, *μ*_2_) can be calculated as:

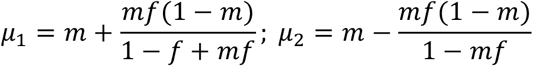 The hyper-prior on *f* is modeled by a uniform beta distribution, which means the fraction of abnormal cells ranges from 0% to 100% with equal prior probability before inference:

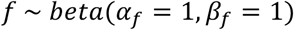 Thus, under this model, the likelihood of the data is given by:

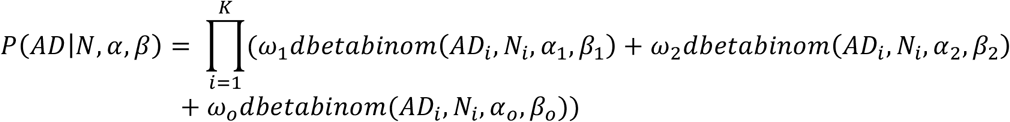
3. **Model_2_: mosaic mitotic trisomy.** Mosaic mitotic trisomy arises from non-disjunction during mitotic cell division, resulting in an extra copy of one of the normal chromosomes. As a result, the AAF at heterozygous sites will be separated into 2 mixtures, in a qualitatively similar pattern to that of mosaic monosomy case (**Figure S2**). However, the expected average AAF of the mixtures for a given fraction of aneuploidy differ, with the means expected to be given by:

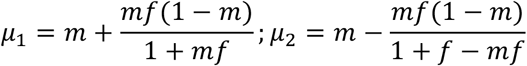 The hyperprior on *f* and the weights for separated mixtures *ω*_1_ and *ω*_2_ and the outlier component are the same as described in mosaic monosomy.
4. **Model_3_: mosaic meiotic trisomy.** Trisomy can also be acquired during meiotic cell division. Mosaic meiotic trisomy can be distinguished from mitotic trisomy by the presence of two additional mixtures near the boundaries (**Figure S2**), which are the consequence of recombination during meiosis. Based on the assumption that the four separated mixtures have the same variance (*κ*), the allelic depth of heterozygous sites can be modeled as:

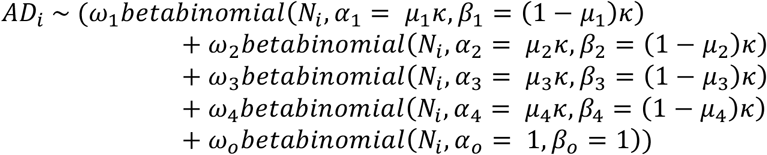 Among the four separated mixtures, the two mixtures in the center are expected to have the same means as described in model_2_ (*μ*_1_, *μ*_2_). The means of the two additional mixtures near the boundaries are given by:

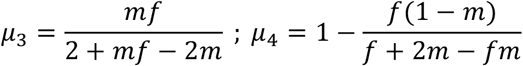 We assume that the two central mixtures have the same weight, and that the two edge mixtures also have the same proportion. As the edge mixtures can have smaller or equal weight compared to the center mixtures, we therefore model the prior of the weight of the edge mixtures by a truncated uniform beta distribution:

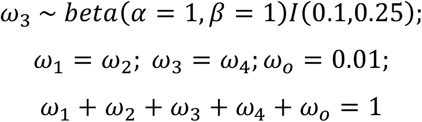 The hyper-prior *f* is modeled the same way as described above.
5. **Model**4**: mosaic loss of heterozygosity (LOH).** Mosaic copy neutral loss of heterozygosity can be due to multiple reasons, for example due to trisomy rescue when the whole chromosome is involved, or recombination-mediated repair when the LOH is segmental. As a result, the AAF of the heterozygous sites or all or some of the chromosome will also be split into two mixtures (**Figure S2**). To characterize such regional effects of LOH, we introduced a reversible jump model, which contains two change points (*cgp_s_, cgp_e_*) to account for the start and end of the LOH status, to describe the combination of normal and abnormal regions on the same chromosome. In the normal regions, the model is the same as for a normal chromosome:

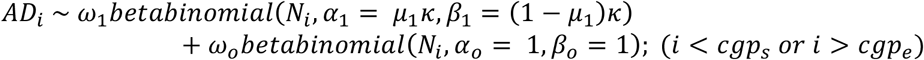 In the LOH region, the distribution of AAFs is separated into two mixtures:

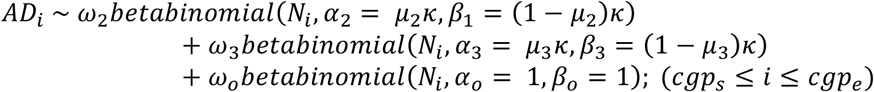 For the normal region (*i* < *cgp_s_ or i* > *cgp_e_*), the distribution is modelled in the same way as for a normal chromosome, with *μ*_1_ calculated here as the average AAF for heterozygous sites. For the LOH region, the weights for the two mixtures are assumed to be the same. The means of the separated mixtures are calculated as:

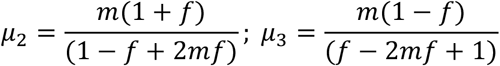 The hyper-prior *f* is modeled the same way as described above. Given there is a total of *K* heterozygous sites on each chromosome, the priors of the changing points, which are the starting locus and ending locus of the abnormal region, are modelled by two uniform distributions ranging from 1 to *K*. In order to be robust against noise in the data, we require that the LOH region spans at least 10% of the total number of loci (*K*) on one chromosome:

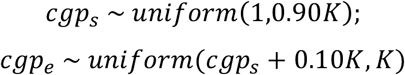

**Figure 1.**
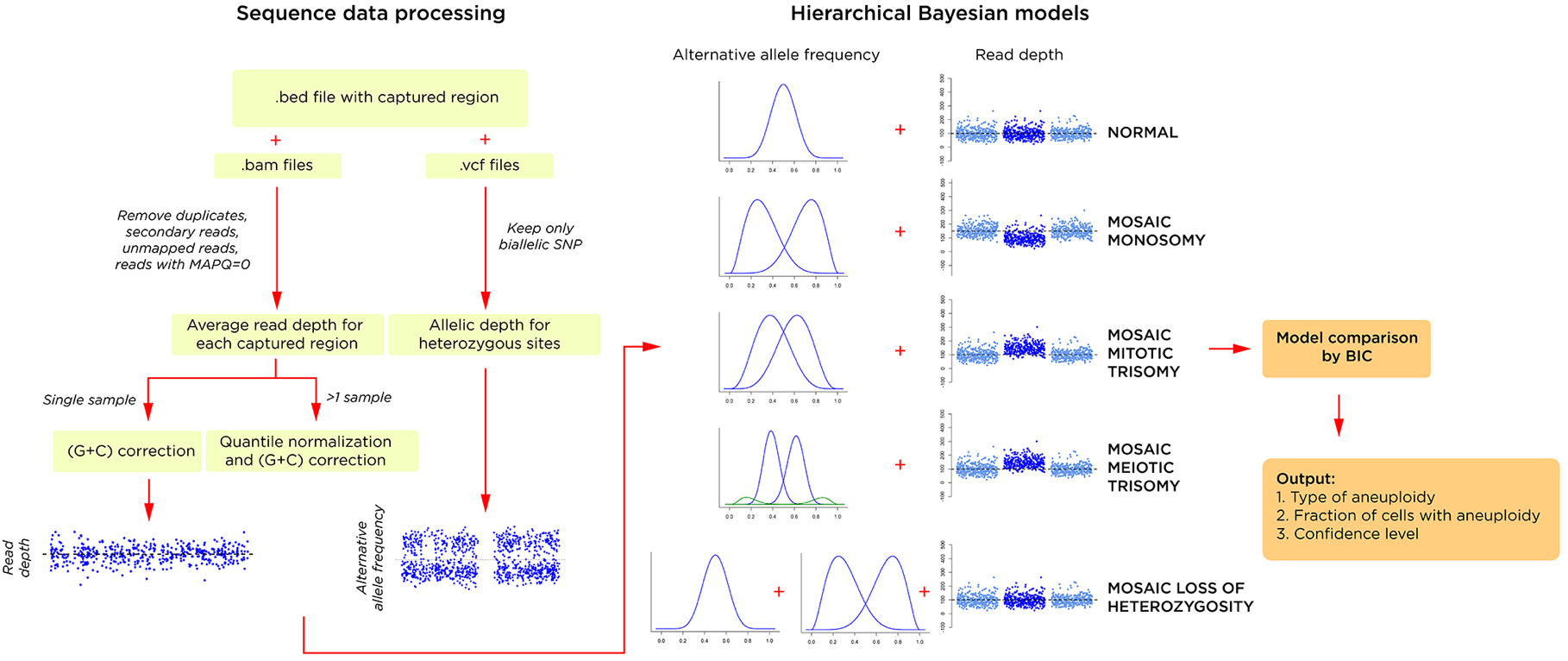
The *MADSEQ* analytical approach has two major components, the processing of the sequence data (left) and the use of alternative allele distributions and relative read depth to support a specific hierarchical Bayesian model (right). The winning model is selected by its significantly better Bayesian information criterion (BIC), generating an output of the presence and type of mosaic aneuploidy (or loss of heterozygosity), the proportion of cells with the aneuploidy, and the confidence of this prediction based on relative BIC values.

**Figure 2.**
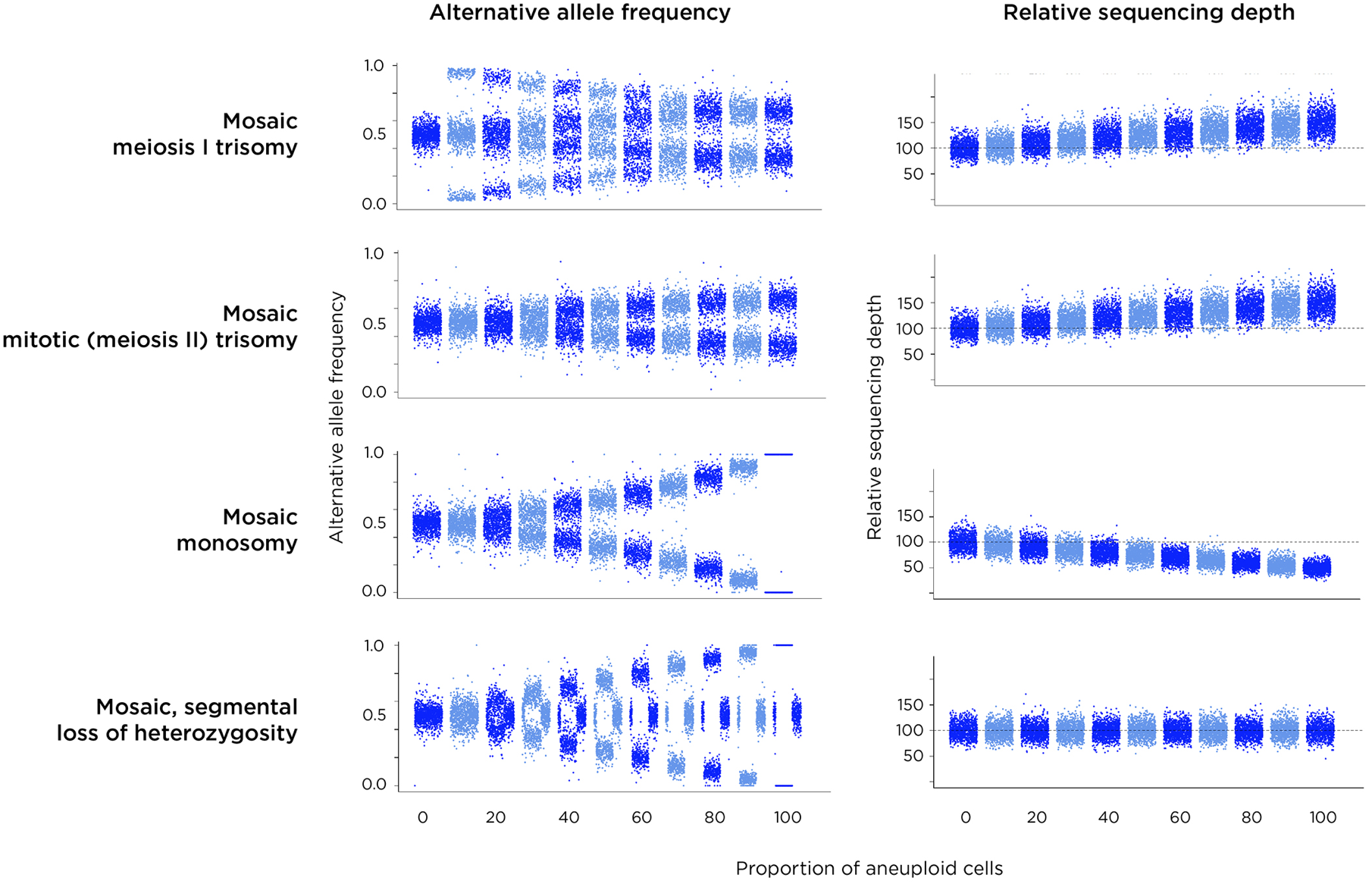
Results of simulations of each of the types of aneuploidy (plus loss of heterozygosity, LOH). Each plot on the **left** shows the AAF distributions in steps of 10% proceeding from no aneuploidy (complete diploidy) to aneuploidy in all cells. On the **right** is shown the expected change of sequencing depth for the affected chromosome. Because meiosis I involves the third chromosome in the mosaic trisomic cells having a different haplotype to those in diploid cells with which they are mixed, the AAF pattern is very distinctive (**top**), with coverage for the chromosome increased relative to others in the genome. Trisomy occurring in meiosis II or mitosis should look We show simulated results for four types of chromosomal mutations in **Figure S2**, monosomy, mitotic and meiotic trisomy, as well as copy neutral loss of heterozygosity (LOH). We include LOH as it could represent in diploid cells the consequence of trisomy rescue earlier in development^26^, an aneuploidy-related event. However, we developed the approach so that it could also detect segmental LOH occurring in >10% of contiguous heterozygous sites tested in each chromosome. What is apparent from **Figure S2** is that our ability to detect and discriminate the different types of aneuploidy events depends upon the alternative allele frequencies in combination with any deviation from the genome-wide average coverage of sequence reads for that chromosome. For example, while mosaic monosomy and mosaic mitotic trisomy have similar alternative allele frequency patterns, they differ by coverage, with trisomy generating an excess and monosomy a deficiency of sequence reads for that chromosome compared with the remainder of the genome.

##### 2. Inference of the type of aneuploidy from sequencing depth

While the distribution of AAF is informative of mosaic aneuploidy, it is difficult to distinguish between the cases without additional information, as, for example, mitotic trisomy and mosaic monosomy have similar distribution of AAFs. In order to improve our differentiation of different types of aneuploidy, we augmented our model with information about sequencing depth.

In our model, the expected coverage for normal chromosome is denoted by *m_g_*. If there is only one sample, is calculated as the median of GC corrected coverage for the whole genome. If there are multiple samples, *m_g_* is calculated as the median across the normalized coverage for that chromosome across all samples.

For the sequencing depth, the total number of targeted regions from one chromosome is *nRegion*; the coverage of the *i^th^* region is *cov_i_*. In order to characterize the over-dispersion of the depth observed in massively-parallel sequencing data, we model the coverage as a negative binomial distribution:

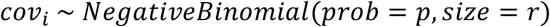

where the prior of *r* is modeled by a weakly informative gamma distribution:

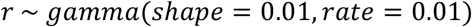

and *p* is taken as:

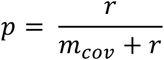

Here, *m_cov_*, which is the mean of the coverage for this chromosome, is determined by the expected normal coverage *m_g_* and the fraction of aneuploid cells *f*:

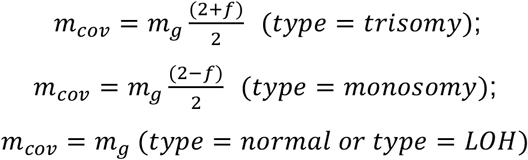

In this way, we could further optimize the estimation for the fraction *f* through the coverage information, while at the same time better inferring the type of aneuploidy. The likelihood of the coverage data over all sites is:

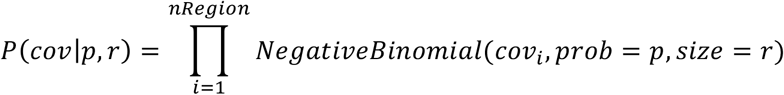

To combine information from the AAF and coverage models, we take the combined likelihood as:

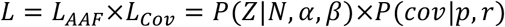

#### MCMC sampling

Have the likelihood function and prior set for each model, the posterior distribution is sampled through Markov Chain Monte Carlo (MCMC). The sampling process is done using JAGS in the R package *rjags*. The script for the model is included in the *MADSEQ* package at Bioconductor. For all the sample and computational simulation, we set the burn-in steps to 10,000, and we sampled two chains, each with total 10,000 steps and each step sampled at every 2 steps. The convergence of the two chains is checked using the Gelman and Rubin diagnostic with the *coda* package in R.

#### Model comparison

After we get the posterior distribution from MCMC sampling, the goodness of fit of models are compared using the Bayesian information criterion (BIC). The exact maximum likelihood of each model cannot be calculated directly from the MCMC procedure because of the hierarchical nature of the model, so we take a point estimate of the likelihood for each model using the median of each parameter from the posterior distribution^37^.

Ultimately, the model with the lowest BIC is preferred as the best model, and the type and fraction of aneuploidy are estimated from the posterior distribution of the best model. If the ΔBIC between the selected model and other models is less than 10, then we consider the chosen model to be low confidence^15^.

#### Computational simulations

We aimed to evaluate the performance of our model as a function of sequencing depth, the type and fraction of aneuploidy cells, and the number of heterozygous sites sequenced for one chromosome. We randomly generate data as follows:

1. **Simulation of coverage.** Given the expected normal coverage m_cov_ and the fraction of the aneuploid cells *f*, the average coverage for the chromosome *m_cov_* can be calculated as described above. The sequencing depth *cov_i_* for the *i^th^* site was randomly drawn from the negative binomial distribution:

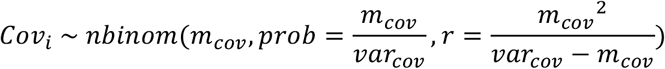 We set the variance of the coverage (*var_cov_*) to 30 times of *m_cov_* based on what we observed from the actual sequencing data. The total number of targeted regions (*nRegions*) was fixed as 1.5 times of the total number of heterozygous sites (*K*).
2. **Simulation of AAFs.** Given the type of aneuploidy, the fraction of abnormal cells (*f*), the total number of heterozygous sites (*K*) on one chromosome and the midpoint AAF (*m*) across all heterozygous sites. The mean AAF for each mixture (*μ_j_*) is easily calculated using the formula described in the model section. The weight for each mixture (*ω_j_*) is given by the same way as in the model, we randomly assigned *ω_j_*K sites into the *j^th^* mixture. Knowing the average coverage for the simulated chromosome (*m_cov_*), the read depth for the *i^th^* heterozygous site (*N_i_*) is also random drawn from the negative binomial distribution:

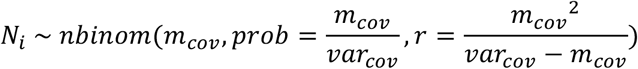 The alternative allelic depth for the *i^th^* site (*AD_i_*) is randomly generated from the binomial distribution as:

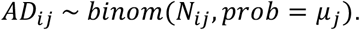
3. **Simulation of noise.** To account for the influence of noise in real sequencing data, we randomly selected 1% of the sites to have an alternative allele frequency drawn from a uniform beta distribution. We also randomly picked 1% of the regions to have random coverage uniformly spanning from 1 read to the maximum amount of coverage. When testing the false positive rate, we increased the noise level to 10% instead.

Since we only use sites that are genotyped as heterozygous to estimate the distribution of AAFs, we have to consider the capacity of the genotyping algorithms to call heterozygous sites. In general, genotyping algorithms will call a site as heterozygous if there are multiple reads supporting each allele. We therefore filtered out sites with fewer than 3 reads supporting both the alternative and reference alleles, and sites whose AAF are less than 0.02 or greater than 0.98 from the simulated data. We simulated 500 sets of data for each aneuploidy scenario.

#### Exome sequencing data from the 1000 Genomes Project

The BAM files of exome sequencing data of 2,535 individuals from 1000 Genome Project were downloaded from the FTP site of the 1000 Genome Project: ftp://ftp.1000genomes.ebi.ac.uk/. The BED file containing the targets of the exome sequencing was downloaded from: ftp://ftp.1000genomes.ebi.ac.uk/vol1/ftp/technical/reference/exome_pull_down_targets/

#### Design of multi-ethnic targeted loci

We show the steps involved in **Figure S7**. Genotyping data of 1000 Genomes Project were downloaded from: http://ftp.1000genomes.ebi.ac.uk/vol1/ftp/release/20130502/

First, we kept only biallelic loci. The heterozygosity rate for each locus was then calculated as:

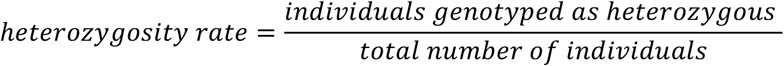

Loci with a heterozygosity rate greater than 0.4 were retained. Loci located in repetitive regions annotated by the RepeatMasker (rmsk) track from the UCSC genome browser were excluded. Loci located within ±20 bp of known indels and within ±500 bp of gaps were excluded. In order to decrease (G+C) content bias, we removed loci located within extreme (G+C) content regions ((G+C)<0.3 or (G+C)>0.65, 200 bp context).

According to the computational simulation, the model can achieve very high sensitivity when there are ≥2,000 heterozygous sites sequenced on each chromosome. As the mean heterozygosity for the loci we retained was 0.45, we aimed to keep 5,000 loci per chromosome for the targeted sequencing.

In order to make loci evenly distributed along the chromosome instead of forming clusters, we binned each chromosome into 500 equal sized windows using bedtools^38^. We then randomly selected ~10 loci from each window. In total, we created a list containing 105,703 common SNPs for further capture. We used a Q-Q plot to show that there was no clustering of loci compared to randomly selected loci.

#### Alignment, genotyping and data processing

Raw FASTQ files from the sequencing were aligned to the GRCh37 human reference genome using *BWA-MEM* (v0.7.10) on default paired-end mode^26^. *Picard* (v 1.119) was used to mark duplicates and *GATK* (v 3.4-46)^25^ was used for indel realignment and base recalibration following best practices^27–29^. *HaplotypeCaller* was used to call variants and to genotype all of the targeted sites.

#### Estimation of ancestries of samples

We performed principal component analysis (PCA) using *EIGENSTRAT*^39^ across 12 samples sequenced by meMAD-seq assay and 2,054 samples (26 populations) in the 1000 Genomes Project. SNPs used for PCA analysis are 1000G SNPs located in the captured regions of the meMAD-seq assay. *Vcftools*^40^ is used to process the SNPs.

#### *MADSEQ* model application

Having aligned the BAM file, genotyped the VCF file and prepared the BED file containing the targeted regions, we used the *MADSEQ* package to correct for GC bias and filter noise, running the *MADSEQ* model as described in the documentation on Bioconductor.

#### Statistical analysis

We performed binomial test and chi-square test to test enrichment and association between detected aneuploidy and other factors. All the statistical testing was performed in R (v3.2)^41^.

## RESULTS

### The MAD-seq molecular assay

In a single sample of cells, chromosome aneuploidy is revealed by altered dosages of minor alleles, usually referred to as alternative allele frequencies, throughout a chromosome. For example, normally an alternative (B) allele can be present at a diploid locus at 0% (AA), 50% (AB) or 100% (BB) frequencies, but a cell with a chromosome trisomy will have 0% (AAA), 33% (AAB), 67% (ABB) or 100% (BBB) frequencies for that chromosome. Mosaicism for the trisomy in a population of otherwise diploid cells is reflected by values intermediate between these extremes, with the proportion of mosaic cells reflected by the relative extent to which the minor allele frequencies resemble the diploid pattern (indicating low-level mosaicism) or the trisomic pattern (indicating high-level mosaicism). With greater numbers of loci representing each chromosome, and more of these loci heterozygous in the individual tested, there will be greater ability to detect and quantify mosaic aneuploidy in a cell sample from that person.

We therefore designed a trial customized SeqCap (Roche-NimbleGen) assay (v1MAD-seq) capturing 80,000 loci in the human genome, targeting loci with highly polymorphic single nucleotide polymorphisms (SNPs). These SNPs were selected based on being represented on the Illumina HumanOmni2.5 genotyping array and having been studied by the HapMap project, and by each having a minor allele population frequency of at least 0.4. The captured loci were distributed evenly in the genome, using capture oligonucleotides designed by Roche-Nimblegen, with most loci having 1-3 tiling probes for a 125 base pair window centered around the interrogated SNP. 79,605 SNPs were included in the final probe capture design.

We performed an experiment using cell-line derived DNA samples from a male Yoruban (GM19239, Coriell Cell Repository) and a female Caucasian (GM06990), mixing the samples to create serial dilutions of GM19239 in GM06990 as 50%, 25%, 10%, 5%, and 0.5% proportions (**Table S1**). A separate sample of GM06990 on its own was prepared as a control (0%). Capture and Illumina sequencing were followed by alignment and processing using BWA (version 0.7.10)^24^, and elimination of PCR duplicates using Picard (version 1.119, https://broadinstitute.github.io/picard/). Variant calling was performed using the Genome Analysis Toolkit (version 3.4-46)^25^, including base recalibration and variant calling using the HaplotypeCaller. We plotted the dilutions of GM19239 in GM06990 DNA against the proportion of GM19239 to GM06990 alleles in **Figure S1a**, showing that the subset of GM19239 reads is clearly detectable down to 5%. We therefore proceeded to develop further an analytical approach that would allow us to detect single chromosome aneuploidy events following such capture.

### The *MADSEQ* analytical approach

We provide an overview of the analytical approach in Figure 1. There are two main components to *MADSEQ*, the processing of the sequencing data and the generation and comparison of hierarchical Bayesian models. The output of the *MADSEQ* analysis consists of (a) the identification of aneuploidy for one or more chromosomes, (b) categorization of the type of aneuploidy, (c) quantification of the fraction of cells with the aneuploidy, and (d) a confidence metric in the results obtained.

We show simulated results for four types of chromosomal mutations detected by the *MADSEQ* analytical approach in Figure 2, monosomy, mitotic and meiotic trisomy, as well as copy neutral loss of heterozygosity (LOH). We include LOH as it could represent the consequence of trisomy rescue earlier in development^26^ in currently-diploid cells. However, we developed the *MADSEQ* approach so that it could also detect segmental LOH occurring in >10% of contiguous heterozygous sites tested in each chromosome. What is apparent from Figure 2 is that our ability to detect and discriminate the different types of aneuploidy events depends upon the alternative allele frequencies in combination with any deviation from the genome-wide average coverage of sequence reads for that chromosome. For example, while mosaic monosomy and mosaic mitotic trisomy have similar alternative allele frequency patterns, they differ by coverage, with trisomy generating an excess and monosomy a deficiency of sequence reads for that chromosome compared with the remainder of the genome. A meiosis II non-disjunction causing trisomy will appear similar to trisomy caused by mitotic non-disjunction, but can be distinguished by the presence of a chromosomal region that underwent recombination earlier in meiosis, which is flagged in the *MADSEQ* analysis when choosing the optimal model.

### Evaluation of *MADSEQ* performance

We tested the performance of *MADSEQ* in two ways. The first evaluation was a re-analysis of the GM19239/GM06990 mixing experimental data, the second based on computational simulations. For the Yoruban/Caucasian mixing experiment, we used a beta distribution to fit the alternative allele frequency (AAF) and measured the deviation in these samples from an expected distribution in diploid cells. We measured this deviation for each chromosome in each of the samples and plotted the relationship between this distance and the expected proportions of GM19239 DNA, as shown in **Figure S1b**. We showed the expected correlation with the known mixture proportions, but this time using our AAF deviation distance, indicating that the model was able to reproduce the information from known Yoruban and Caucasian genotypic differences.

We went on to explore the sensitivity, specificity and accuracy of the *MADSEQ* approach using computational simulations. To assess sensitivity, the performance of the model was tested in terms of sequencing depth, proportion of aneuploid cells present, and the number of heterozygous sites sequenced per chromosome. For example, if 2,000 heterozygous sites on a chromosome are sequenced to 100X coverage, our model should be able to detect 5% mosaicism for meiotic trisomy with greater than 99% sensitivity at a false discovery rate (FDR) of <2% (**Figure S2**). The power to detect low proportions of mosaic aneuploidy increases with deeper sequencing and higher numbers of heterozygous sites sequenced. When 2,000 heterozygous sites are sequenced to 200X coverage, we predict more than 50% power to detect all types of mosaic aneuploidy occurring in 10% of cells. Specificity issues and the generation of false positive results are important for an assay that might be used for clinical diagnostic purposes. We evaluated the FDR of our method by simulating data without aneuploidy but with the simulated introduction of noise in the sequencing data, randomly distributing the alternative allele frequencies of 10% of the heterozygous sites between 0 and 1. The result (**Figure S3**) suggests that our model is very robust with an overall FDR at <2%. The accuracy of the quantification of the fraction of abnormal cells was assessed using the root-mean-square-error (RMSE) calculated from computational simulations. In **Figure S4** we show the quantification is accurate with deviation from the expected proportion of less than 10% for all the conditions tested, with accuracy increasing with deeper coverage but with less effect from including more sequenced sites per chromosome.

### Application of *MADSEQ* to sequencing data

We then explored whether analysis using *MADSEQ* could identify mosaic aneuploidies from publicly-available sequencing data. The 1000 Genomes exome sequencing was generated to a mean of 65.7X coverage, potentially capable of discovering at least some mosaic aneuploidy events if present in these samples. Of the 2,535 individuals studied in the 1000 Genomes Project, 2,037 were sequenced using DNA derived from LCLs, whereas the remaining 498 were sequenced from peripheral blood leukocyte DNA. *MADSEQ* detected 83 mosaic events with high confidence (ΔBIC > 10) in 76 individuals (**Table S2**). All of the detected mosaic aneuploidies were from LCL samples, and none from blood. The types of mosaic aneuploidy include 20 monosomies (0.79%), 25 mitotic trisomies (0.99%) and 37 LOH (1.46%), but no meiotic trisomies, even though the model is relatively more sensitive when detecting meiotic events. Of note, all of the cases of LOH were segmental rather than involving the whole chromosome, likely to represent the result of a repair of deletion using the remaining homologous chromosome as a template^27^, and not trisomy rescue.

The rate of these mosaic events among LCL samples was 3.73%. The proportions of aneuploid/segmental LOH cells in each sample varied from 4.2% to 79.9%. The most overrepresented events were mosaic mitotic trisomy of chromosome 12 in 11 samples (p= 1.49x10^−4^, a Binomial Test), and enrichment for mosaic segmental LOH in chromosome 22 (p= 7.14x10^−3^, Binomial Test) (**Table S3**). Overall, the significant lack of mosaic aneuploidy events in samples from blood (p= 3.08x10^−34^, Binomial Test) and the lack of meiotic events, together with the enrichment of trisomy 12, which is the most common cytogenetic abnormality in chronic B lymphocytic leukemia^28^, combine to suggest that these mosaic aneuploidies arose during cell culture and were either neutral in effect or promoted positive selection for these transformed B lymphocytes. One of the samples in which segmental LOH for distal chromosome 11 was identified was GM12889, for which whole genome sequencing (WGS) to a mean ~50X coverage has been performed to define high-confidence, “platinum” variants^29^. We downloaded those WGS data and re-ran *MADSEQ*, again predicting the LOH of a 19.3 Mb region, estimated to be present in over 50% of the cells (Figure 2). The platinum variant calling in this part of the genome in this individual should be interpreted with caution. We show representative examples of plots of the alternative allele frequencies and the comparisons of the Bayesian Information Content (ΔBIC) in **Figure S5**.

We then tested a sample from a patient presenting with hemihyperplasia (OMIM 235000). The hyperplastic side of the patient demonstrated a hyperpigmented whorl pattern, following Blaschko’s lines. The Blaschko’s line were only present on the hyperplastic side and did not cross the midline. Skin biopsies were performed on the normal skin of the unaffected side, and from the hyperpigmented skin of the affected side. A microarray study from DNA directly extracted from these biopsies showed evidence for mosaic trisomy 12 from the affected side only. We grew fibroblasts from the remainder of the skin biopsy and extracted DNA from these cultured cells, performing exome sequencing to mean 130X coverage. The *MADSEQ* model best fit by the results was of mosaic trisomy of mitotic origin present in 6.8% of the cells (ΔBIC = 18) (Figure 3). Trisomy 12 has also been found in human embryonic stem (ES) cell lines^30^ and induced pluripotent stem (iPS) cells^31^, and has been implicated in the significant increase of cellular proliferation rate and tumorigenicity^20^. The hemihyperplasia phenotype may therefore be the consequence of a higher cell replication rate due to the presence of the mosaic subset of cells with trisomy 12. Our categorization that this was a mosaic trisomy of mitotic origin (Figure 3) indicates that the non-disjunction and chromosomal loss events occurred post-fertilization, and not due to meiotic trisomy during gametogenesis with later trisomy rescue.

**Figure 3.**
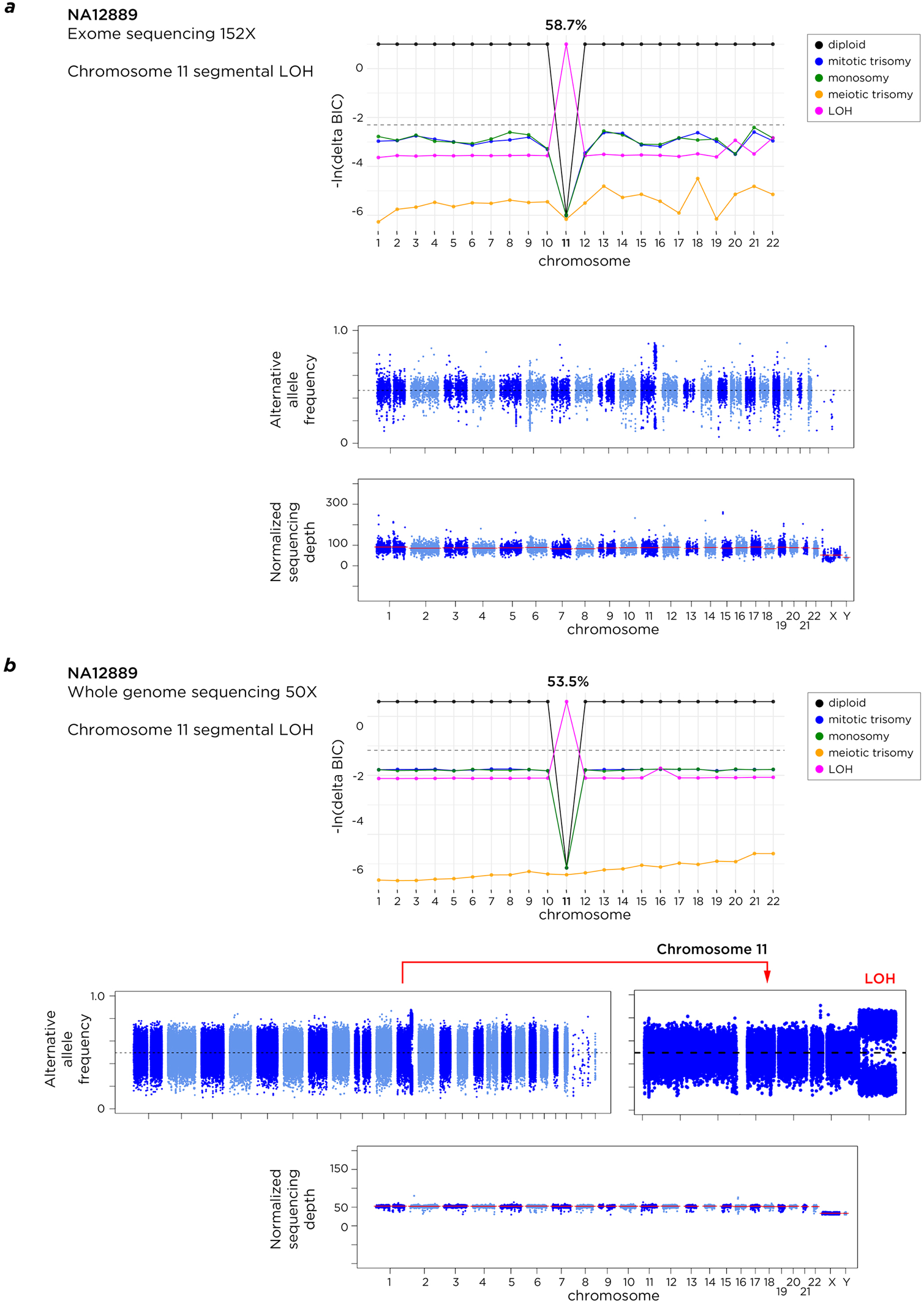
The NA12889 sample has been tested using exome sequencing as part of the 1000 Genomes study (top) and using whole genome sequencing to mean 50X coverage (bottom). The application of the *MADSEQ* approach shows concordant predictions of mosaicism for segmental, copy number neutral LOH in distal chromosome 11 (bottom) in ~54-59% of cells.

### Development of a multi-ethnic MAD-seq (meMAD-seq) assay design

As the depth of exome sequencing (130X) used for sensitive detection of the mosaic mitotic trisomy 12 in only 6.8% of the cells was impractical for a routine test, we returned to our v1MAD-seq capture design and tested its performance with six samples, four of which were known to have autosomal trisomies, and two control cell lines apparently lacking aneuploidies. Using *MADSEQ* analysis of the results, we confirmed the four trisomies (chromosomes 8, 13, 15 or 18), mostly concordant with prior reported proportions of trisomic cells, and defined their origins as meiotic in all cases. One of the control cell lines (GM06990) that had not previously been described to have aneuploidy was found to have a pattern consistent with 6.6% of cells having monosomy for chromosome 6 (Table 1).

**Table 1.**
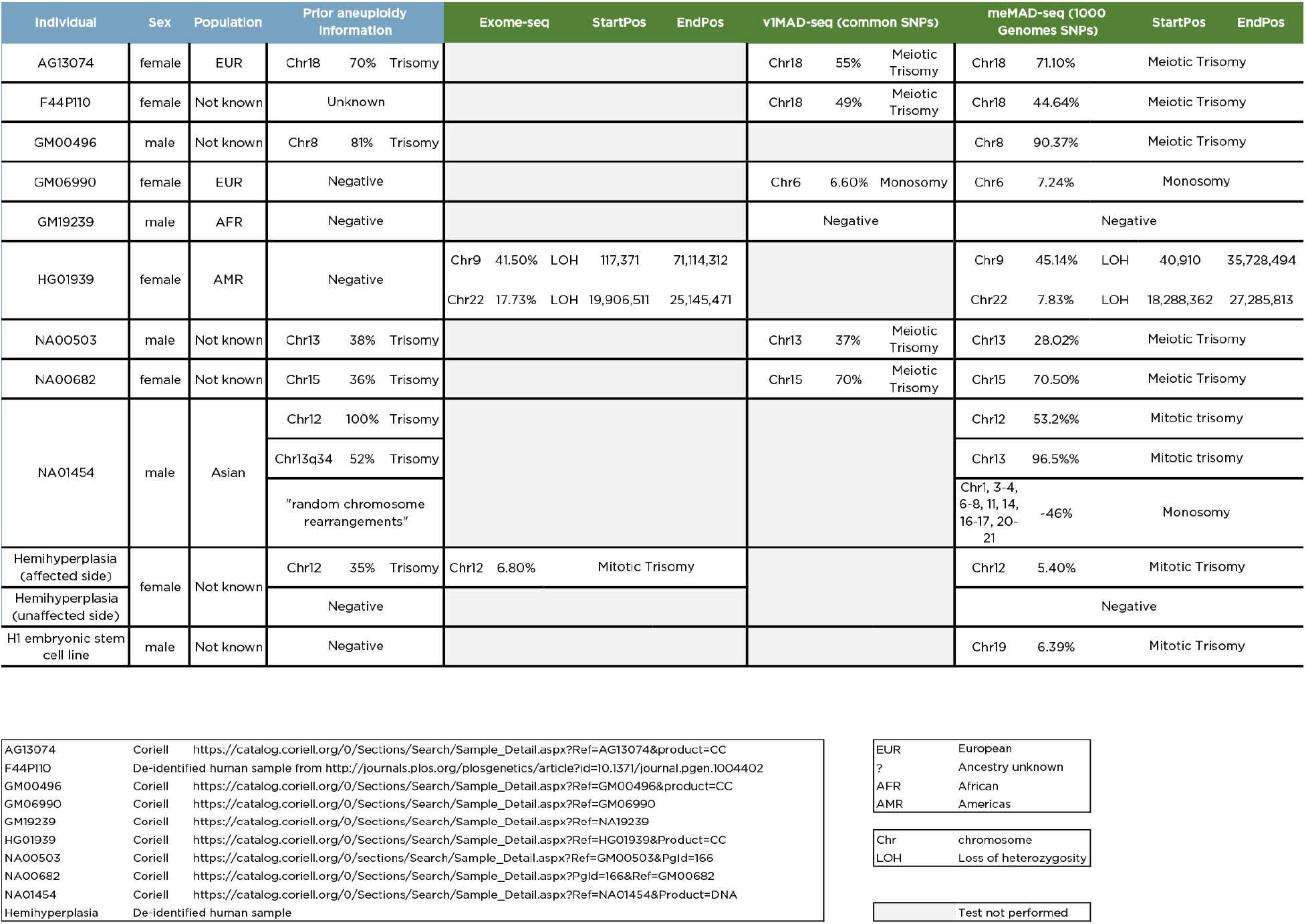
Overview of results from exome sequencing, v1MAD-seq and meMAD-seq assays, compared with information about samples known before testing. We see that in general when there was a known mosaic aneuploidy in the sample, we confirm it, but some samples (HG01939 and H1 embryonic stem cells) that were supposedly negative are found to have detectable mosaic aneuploidies. The proportions of cells with aneuploidy are generally but not always concordant with those identified using our analyses, which also identify the likely mechanism of the aneuploidy.

When we explored the performance of the v1MAD-seq and exome sequencing systems, we found that the representation of heterozygous sites on many of the smaller chromosomes was highly suboptimal. The v1MAD-seq design spaced loci for capture evenly throughout the genome, causing larger chromosomes to have proportionately more informative loci (Figure 4a) while exome-seq is limited by the number of genes per chromosome, which is a function of not only the chromosome size but also its gene content. In Figure 4b we show this heterogeneity of representation for each chromosome for exome sequencing data. Of particular concern was the poor representation of informative loci for chromosomes 13, 18 and 21, the most common viable full trisomies.

**Figure 4.**
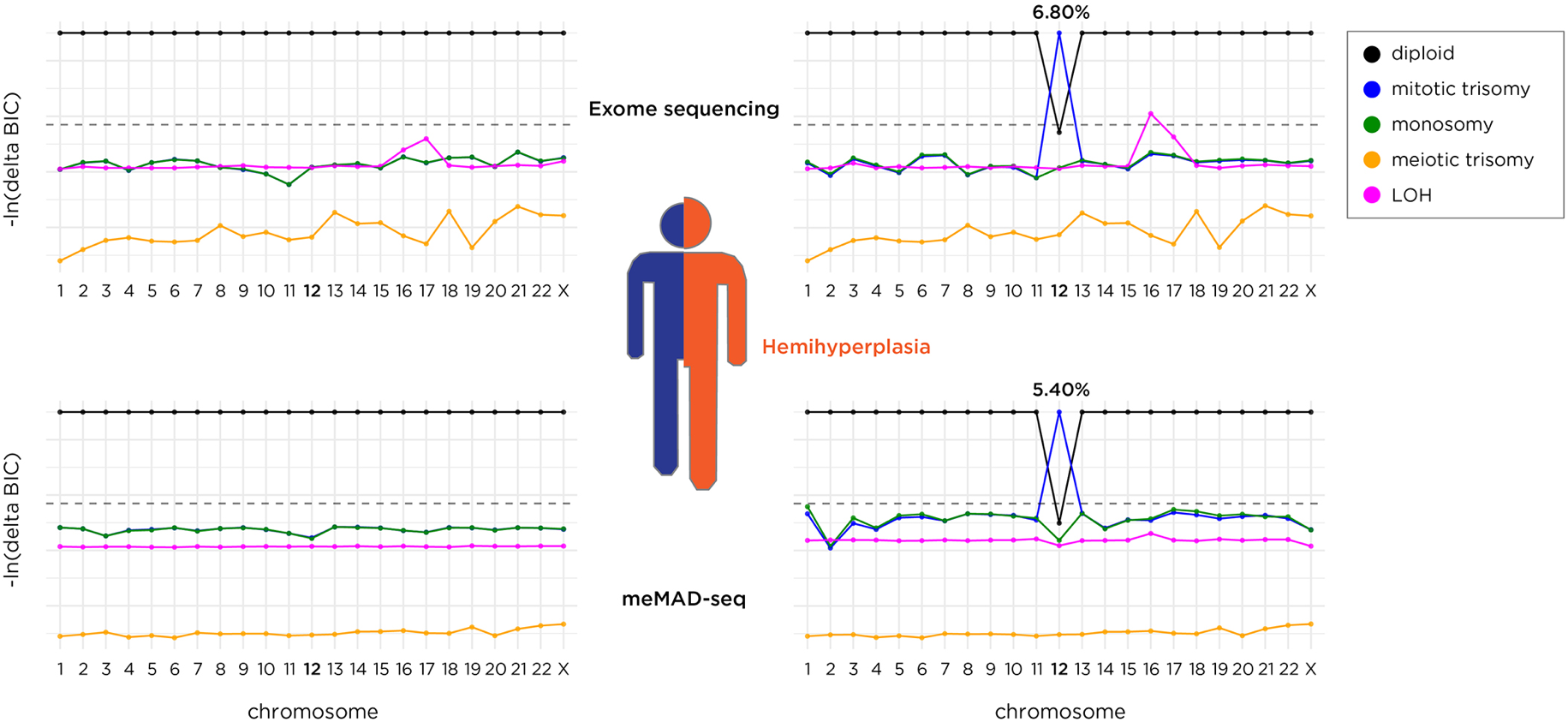
The results of sequencing skin biopsy samples from each side of the body of a child with hemihyperplasia. Results from the normal side are shown on the left, and from the overgrown side on the right. Exome sequencing to ~100X mean coverage favors a model of mosaic trisomy 12 of mitotic origin in 6.8% of cells, while the more targeted meMAD-seq assay sequenced to a comparable depth shows evidence for the same type of mosaicism in 5.4% of cells.

To create a design that is maximally efficient in sequencing informative loci, with resulting efficiencies in assay cost, we created a new multi-ethnic MAD-seq (meMAD-seq) assay design. At total of 107,797 (Roche NimbleGen Catalog No. 06740260001, Design Identifier: 160407_HG19_MadSeq_EZ_HX1) common SNPs were chosen to represent each chromosome equally. We also exploited 1000 Genomes data to identify loci that would be most likely to be polymorphic across all human populations (**Figure S6**). We show the workflow for the design of the meMAD-seq platform in **Figure S7**. This design captures 106,402 loci in the human genome of mean length 139 bp and mean (G+C) content 44.4%.

We tested the meMAD-seq design on 12 samples. These included the 6 samples tested using the v1MAD-seq design, the HG01939 sample predicted to have two separate chromosomes with loss of heterozygosity from our 1000 Genomes exome sequencing re-analysis, samples from the affected and unaffected sides of the body of the individual with hemihyperplasia, and two Coriell cell repository samples, one of which was described to have mosaic trisomy 8 (GM00496), and a sample described to have mosaic trisomy 12 as well as random chromosome rearrangements (NA01454), and DNA from the H1 human embryonic stem (ES) cell line^32^. We show the results in Table 1 and Figure 5, confirming prior observations of chromosomal aneuploidies or loss of heterozygosity from Coriell’s characterizations, our exome sequencing data re-analysis, or the v1MAD-seq results, and adding information about whether each trisomy was likely to be meiotic or mitotic in origin.

**Figure 5.**
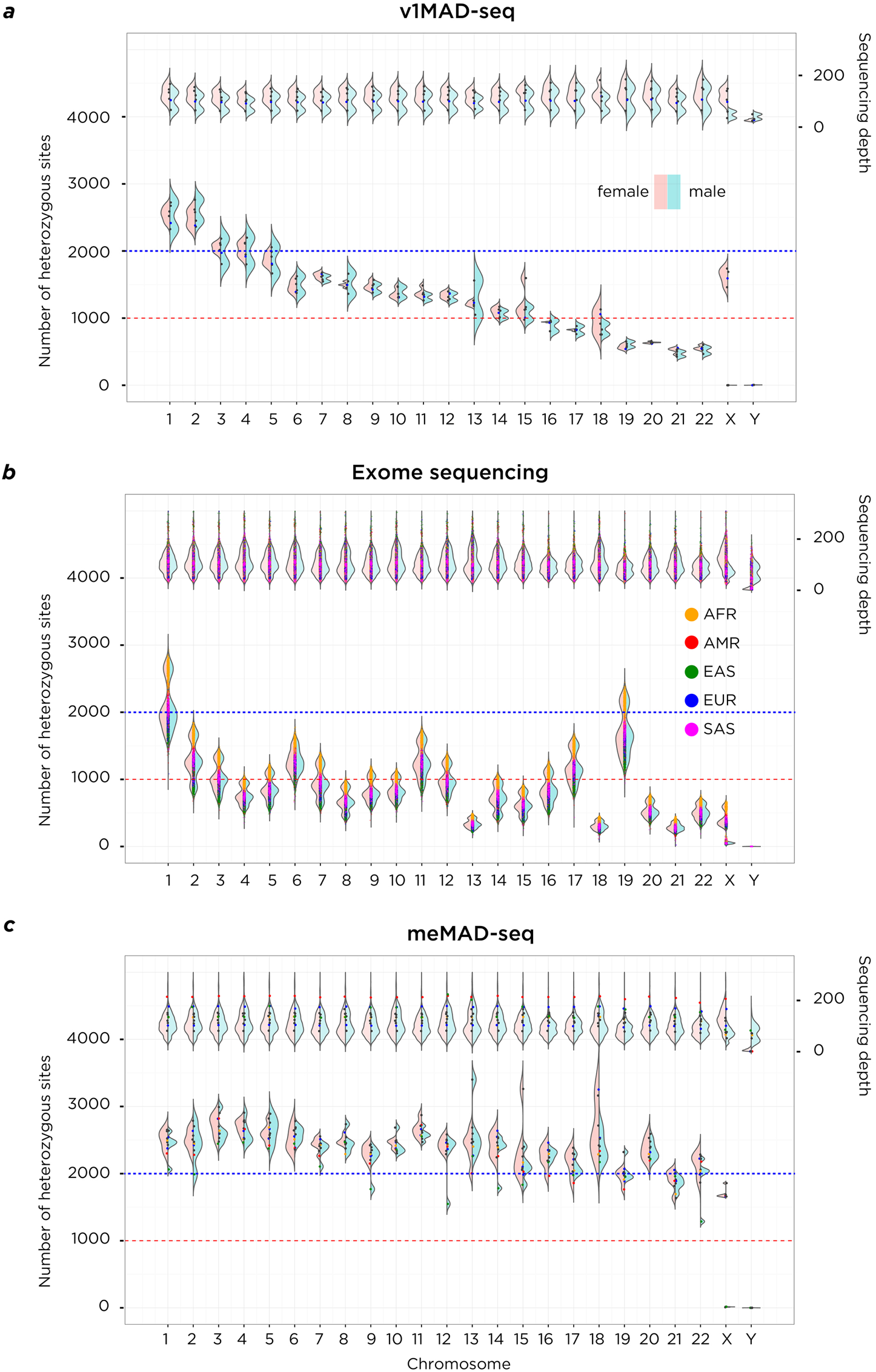
Performance comparison of v1MAD-seq, exome sequencing and meMAD-seq in terms of the number of heterozygous sites tested per chromosome. On the top of each plot is sequencing depth, on the bottom the number of heterozygous sites per chromosome. On the left of each violin plot is the distribution in females (pink), on the right males (red), allowing differentiation of patterns on the sex chromosomes. The v1MAD-seq platform spaced probes evenly throughout the genome, generating more heterozygous sites in larger chromosomes despite equivalent coverage. Exome sequencing (1000 Genomes data) generates a number of heterozygous sites that depends on chromosome size and gene density within the chromosome, so that the comparably-sized chromosomes 18 and 19 differ because of the much greater gene content of the latter. There is also an ancestry effect on heterozygosity rates, with African ancestry performing best and Asian ancestry worst in generating heterozygous sites. The meMAD-seq design allows much more uniform performance of every chromosome in the genome, most exceeding 2,000 heterozygous sites per chromosome.

Our 1000 Genomes SNP selection strategy was designed to generate data from the same number of heterozygous sites for each chromosome and across all populations. Our goal was to exceed 1,000 heterozygous sites per chromosome, but we show in Figure 4c that we obtain ≥2,000 heterozygous loci for every chromosome across all individuals. In **Figure S9** we show that the individuals tested using the meMAD-seq design were indeed from widely-divergent human population groups. We confirmed mosaic aneuploidies in samples known to have these abnormalities from Coriell’s characterization or from our prior studies. Two chromosomes with segmental LOH predicted from our 1000 Genomes exome sequencing data analysis were confirmed in the HG01939 cell line using the meMAD-seq assay. Some of the supposedly normal control samples were also revealed to have aneuploidy, including low level, previously unrecognized events in the ES cells and the GM06990 female Caucasian sample used in our initial serial dilution experiments.

For a cost comparison, we determined the reagent cost expense associated with library preparation, capture and sequencing to comparable depth for the more mainstream exome sequencing approach (SeqCap EZ, Roche-NimbleGen) and the meMAD-seq alternative. We estimate that for each assay to generate mean ~110X coverage, the reagent cost for meMAD-seq would be ~40% of the cost for exome sequencing, which should be a generalizable guide for facilities with different costs. With increased production of the meMAD-seq capture kit, further cost savings may be possible.

## DISCUSSION

This study further strengthens the concern that mosaic aneuploidy is likely to be underrecognised. The 1000 Genomes project performed extensive analysis and quality assessment of their samples and data^33^, but even these carefully studied samples have mosaic aneuploidy in several percent of the LCL samples studied, estimated by *MADSEQ* to involve as many as ~80% of cells. While we interpret the results to indicate that these cases of mosaicism arose during cell culture, this finding should increase the caution required when interpreting information from LCLs in terms of their representation of the donor’s chromosomal status.

We also find that reference cell lines that have been characterized using standard techniques have evidence for chromosomal abnormalities. The human H1 ES cell line has previously been found to develop trisomies *in vitro*^30^, requiring periodic testing to ensure that the cells being used experimentally remain diploid. Reference cell lines supplied by repositories or used in large studies also require careful characterization to ensure that they are not undergoing alterations that could lead to issues of reproducibility of results. It should be stressed that the poor representation of certain chromosomes in exome sequencing data, coupled with some of the 1000 Genomes samples having relatively lower mean coverage, combine to indicate that we are probably missing some further cases of mosaic aneuploidy. The systematic application of meMAD-seq would probably reveal an even higher proportion of reference cell lines with mosaicism.

The analytical software *MADSEQ* is open source, available through Bioconductor, and can be applied not only to meMAD-seq data, but also as we show exome sequencing and even whole genome sequencing data. It could therefore be applied retrospectively to legacy sequence data to look for aneuploidy, generating preliminary results that could then prompt the application of meMAD-seq for a more systematic study. It appears that more development will be needed to extend the use of *MADSEQ* to cancer samples. When multiple chromosomes have abnormal copy numbers, determining what coverage value represents diploidy becomes difficult, weakening a foundational component of the analysis. Our NA01454 sample is not from a cancer, but has multiple chromosomes with abnormal patterns of AAFs and copy numbers, and helps to illustrate how the approach starts to have difficulties when many chromosomes are affected. This will be a focus of further algorithm development, but in the short term *MADSEQ* is valuable for detecting the presence of aneuploidy in even these complex samples. With further development, *MADSEQ* could also be used to detect mosaicism for copy number variants (CNV). However, the resolution of detection will differ based on spacing of the captured heterozygous loci, which in the meMAD-seq design is higher for larger chromosomes, with more physical clustering of loci in smaller chromosomes. The most appropriate future application of *MADSEQ* for mosaic CNV identification may be from WGS data.

Ideally, prospective studies will use the targeted sequencing option of meMAD-seq, allowing optimal cost and performance. The meMAD-seq design, which will be made publicly available through Roche-NimbleGen, shows excellent performance not only in terms of maximising the number of informative sites per chromosome, but also testing each chromosome comparably. We were careful to ensure that the design could be applied equally effectively to people of widely differing ancestries, allowing it to be used world-wide and in our local, highly diverse clinical population.

We anticipate several areas of human disease research that would immediately benefit from MAD-seq. In prenatal genetics care, screening is performed looking for chromosomal aneuploidies, increasingly using non-invasive prenatal testing (NIPT) of cell free DNA in the maternal blood. Positive results from this screening approach can be followed up with invasive tests of fetal cells (chorionic villus sampling, amniocentesis), which can then, in a proportion of cases, lead to discordance between aneuploid NIPT and normal fetal chromosomal results. This situation is presumed to be due to confined placental mosaicism (CPM) for the aneuploidy, but this diagnosis can only be made with the certainty afforded by the sensitivity of the test used on the fetal cells. There is potential for meMAD-seq and *MADSEQ* analysis to enhance the sensitivity of these diagnostic studies. A second potential area worth exploring for covert aneuploidy is in individuals with autism spectrum disorder (ASD). We have noted in a prior study^34^ the association between advanced maternal age and the risk of having a child with ASD^35^. The increased non-disjunction rate in oocytes from older mothers suggests that chromosomal aneuploidy should be tested as a possible mediator of this association, but at present there is little evidence for aneuploidy in individuals with ASD. A more sensitive assay like meMAD-seq applied to samples other than blood from individuals with ASD born to older mothers may be worth exploring as one potential cause of this heterogeneous condition.

The meMAD-seq assay combined with the *MADSEQ* analytical approach can be used on uncultured cells, detects low levels of aneuploidy, identifies the likely mechanism of the initial causative event, is relatively cost efficient, and can be used in any ancestral background. It combines many of the advantages of existing assays to detect aneuploidy and should be suitable for high throughput studies. The eventual goal should be to associate different types of mosaic chromosomal events with human phenotypes.

### Data availability

The sequence data from the F44P110 and hemihyperplasia patients are in the process of submission to dbGaP. The remaining v1MAD-seq and meMAD-seq sequencing data will be available from the Short Read Archive (accession for reviewers at the following link):

ftp://ftp-trace.ncbi.nlm.nih.gov/sra/review/SRP105435_20170501_152247_5d6182b8169f820c3e247e91131138ea

The *MADSEQ* package is available from Bioconductor: http://bioconductor.org/packages/MADSEQ/

The meMAD-seq capture design is available from Roche-NimbleGen: Catalog No. 06740260001, Design Identifier: 160407_HG19_MadSeq_EZ_HX1

## ACKNOWLEDGEMENTS

The authors acknowledge the financial support of the Human Genetics Program, Department of Genetics, Albert Einstein College of Medicine.

## AUTHOR CONTRIBUTIONS

AA, JMG, YK conceived project, led investigation, interpreted results. YK, ERB, AM, KQY, AA developed methodology, processed data, analysed data. ERB, SBM, CAS-P, MS created novel reagents, performed experiments. DFK obtained specimens. YK, ERB, JMG wrote manuscript draft. All authors approved final manuscript.

## MATERIALS AND CORRESPONDENCE

John M. Greally, Department of Genetics and Center for Epigenomics, Albert Einstein College of Medicine, 1301 Morris Park Avenue, Bronx NY 10461, USA.

+1 718 678 1234

